# Adults up to 80 years old maintain effective movement planning when facing complex body dynamics

**DOI:** 10.1101/2025.10.03.680305

**Authors:** Anouck Matthijs, Anda de Witte, Dante Mantini, Jean-Jacques Orban de Xivry

**Affiliations:** Movement Control and Neuroplasticity Research Group, Department of Movement Sciences, KU Leuven, Leuven, Belgium; Leuven Brain Institute, KU Leuven, Leuven Belgium

**Keywords:** aging, feedforward control, feedback control, intersegmental dynamics, shoulder stabilization, electromyography

## Abstract

Aging can significantly impact motor performance, especially in highly complex tasks such as multi-joint movements where the nervous system needs to adequately coordinate mechanical interactions between joints. This coordination is inherently challenging for the brain. Effective coordination of multiple joints relies on intact feedforward control to predict movement dynamics in the initial phase of the movement, and on feedback control to fine-tune the execution in the final phase. However, the effect of aging on these specific control mechanisms remains controversial. In our experiment we investigated a pure elbow motion task using the KINARM exoskeleton. A group of 50 young (20-35 years old), 80 old (55-70 years old) and 30 older-old (80+ years old) healthy participants were recruited. Each participant performed 30° elbow rotations while stabilizing the shoulder joint. Movements were directed toward two distinct targets in both flexion and extension directions. The task was performed under two controlled speed conditions to maximally challenge the motor system, as higher elbow velocities increase interaction torques at the shoulder, demanding greater neuromuscular effort for stabilization. The timing and magnitude of anticipatory EMG activity of the agonist shoulder muscle, necessary to counteract interaction torques, were preserved across all age groups. Moreover, increasing elbow velocity did not result in any performance differences between young and older old adults, indicating that shoulder stabilization during movement initiation remained intact with age. However, older adults exhibited reduced ability to stabilize the shoulder position until the end of the movement, leading to decreased reaching accuracy with older age. These results suggest that feedforward control, essential for movement planning, which is essential for shoulder stabilization during initiation, is preserved during healthy aging and remains resilient to increased motor demands, even in older old adults. In contrast, feedback control appears to deteriorate with age, potentially contributing to reduced movement precision in the final phase of the multi-joint movement.

## Introduction

An essential component of movement control is the coordination of multiple joints to achieve motor goals. Everyday actions such as walking or grasping objects typically require the nervous system to adequately coordinate multiple joints. This coordination is complicated by the complex dynamics of the body, leading to mechanical interactions between limb segments. This means that when a torque is applied at one joint, it will inevitably generate rotational forces at adjacent joints, thereby producing motion even without direct muscle activation at those joints (Hollerbach & Flash, 1981).

To effectively control our movements and accurately anticipate these upcoming interactions torques, intact feedforward control signals from the nervous system are considered essential (A. J. Bastian et al., 1996, 2000; Amy J. Bastian, 2006; Gribble & Ostry, 1999; Hollerbach & Flash, 1981). Feedforward control relies on the knowledge of body dynamics. Such knowledge allows the nervous system to predict the mechanical consequences of the planned movement and to generate appropriate motor commands (Wolpert & Flanagan, 2001). Once a movement is initiated, feedback control mechanisms become increasingly important for maintaining movement accuracy (Scott et al., 2015; Todorov & Jordan, 2002). As sensory information, such as proprioception, vision or touch, becomes available during the movement, the feedback system can update the motor plan and perform online corrections in response to any detected movement error or perturbation (Crevecoeur et al., 2012; Orban de Xivry, 2013; Orban de Xivry & Lefèvre, 2016; Scott et al., 2015; Todorov & Jordan, 2002). While feedforward control enables rapid, anticipatory adjustments to dynamic changes, feedback control operates more slowly and ensures the final precision of the movement. The seamless interplay between these two systems is essential for coordinating multiple joints effectively despite complex body dynamics.

During healthy aging, gradual changes in the neuromuscular system may disrupt the balance between feedforward and feedback control, resulting in less effective coordination of multi-joint movements in older adults (Dutta et al., 2013; Seidler et al., 2002; Verrel et al., 2012). However, several studies have demonstrated that feedforward control is preserved during healthy aging and that older adults maintain effective movement planning (Pai et al., 2003; Sager et al., 2024). For instance, Lee et al. (2007) found no age-related declines in the accuracy of a line drawing task involving shoulder-elbow coordination, despite older adults generated lower muscle forces compared to younger adults. This suggests that while aging can alter inter-joint coordination, feedforward control mechanisms relying on knowledge of inter-segmental dynamics allow the central nervous system to maintain the ability to use it for effective movement control during aging. On the other hand, during healthy aging, increased noise in sensory inputs or delays in sensory processing may impair feedback control, resulting in slower or less precise movement corrections in older adults (Konczak et al., 2012). For example, Dutta et al. (2013) found that coordination deficits (greater hand path variability) in older adults were more pronounced during the second half of a reaching task, potentially indicating an age-related deterioration of feedback processes that are crucial for online adjustments of the movement when the hand approaches a target. However, Verrel et al. (2012) found that older participants were still able to stabilize their arm and maintain endpoint accuracy at a pointing task, which could suggest that feedback control mechanisms may remain intact with older age.

Importantly, when investigating age-related effects on motor control, it is crucial to ensure that the motor system is sufficiently challenged. If a task is not challenging enough, age-related differences may be obscured or underestimated. Introducing a standardized physical stressor in a motor task (see complex system approach: Scheffer et al., 2018) offers an effective method for assessing whether an individuals’ motor performance is resilient to age-related decline. For example, Dutta et al. (2013) demonstrated that older adults experienced greater difficulty coordinating arm movements toward a target when its location was uncertain, compared to when it was fixed. Similarly, Lee et al. (2006) found that when arm movements involved directions with higher inertial resistance, placing greater demands on shoulder-elbow coordination, older adults exhibited reduced force production and slower movement execution compared to younger adults. These findings underscore the importance of sufficiently challenging the motor system to reveal age-related effects.

Given the mixed findings on whether feedforward control is affected by healthy aging, and the possibility that the motor system may not have been sufficiently challenged in previous studies that revealed no age-related effects, the impact of healthy aging on feedforward control remains under debate. Furthermore, because most studies have focused on adults not older than 70-80 years, evidence on individuals over 80 years old is lacking. Yet, feedforward control is essential for effective movement control, particularly for coordinating multiple joints in everyday actions. Gaining further insights into how aging affects the motor system is therefore crucial.

In our study we investigated feedforward control across three age groups of young, old and older-old adults (aged 80+ years), performing a pure-elbow motion task. This task required the participants to maintain shoulder stability during an isolated elbow movement, for which anticipatory activation of the agonist shoulder muscle is essential. This anticipatory activation, driven by feedforward mechanisms, will compensate for the interaction torque at the shoulder, generated by the moving forearm. Thus, the only way to produce a single-joint elbow movement is by applying torques both at the elbow (to rotate the lower arm around the elbow) and shoulder joint (to stabilize the upper arm against the interaction torque) (Almeida et al., 1995; Gribble & Ostry, 1999; Maeda et al., 2017). By increasing movement speed of the forearm, we place greater demands on feedforward control mechanisms to maintain the shoulder joint stable during the initial phase of the movement by amplifying the need for anticipatory motor strategies.

We hypothesize that feedforward control mechanisms are largely preserved during healthy aging, despite the general decline in motor performance (Hunter et al., 2016; Tieland et al., 2018; Wu et al., 2021). However, under increased demands of the motor system, anticipatory muscle activation may become inadequate to fully stabilize the shoulder joint during the initial phase of the movement, particularly in adults aged 80 years and older. Furthermore, we investigate whether the preserved feedforward mechanisms are sufficient to preserve movement accuracy by the end of the movement.

## Methods

### 2.1. Participants

In total, we tested 161 participants for the study. Participants from three age groups were recruited; a group of 50 young adults (YA): 20-35 years old, mean age ± SD = 23.3±2; a group of 80 older adults (OA): 55-70 years old, mean age ± SD = 63.9±4.6; and a group of 31 older-old adults (OOA): 80+ years old, mean age ± SD = 82.4±1.6. Data of one 80+ adult was excluded before data analysis because the participant did not follow the instructions correctly. The participants were recruited as part of a larger cross-sectional study investigating the effects of aging on cerebellar motor function. Accordingly, the required sample size was determined based on the number of participants needed to detect structural changes in the cerebellum across all three age groups (Walhovd et al., 2011). Reported effect sizes (Cohen’s *d*) for such group comparisons are greater than 0.65 (YA vs. OA: 0.98; OA vs. OOA: 0.66; Walhovd et al., 2011). Given that age-related structural decline in the cerebellum is hypothesized to underlie many motor deficits in aging (Bernard & Seidler, 2014; Boisgontier, 2015; Boisgontier & Nougier, 2013; Hogan, 2004; Liang & Carlson, 2020), structural effects of this magnitude were deemed sufficient to justify the detection of functional effects. Power analyses indicated that 50 participants in both the young and older age group and 30 participants in the older-old age group would provide 80% statistical power. To facilitate subsequent longitudinal analyses of the older adult group, the sample size of the older adult group was increased to 80 participants. This adjustment was designed to ensure 80% power to detect an expected small effect (*r* = 0.3) while accommodating an anticipated attrition rate of 20% (Bell et al., 2013).

Participants were screened for right-handedness, according to the Edinburg handedness questionnaire (Oldfield, 1971), and cognitive function based on the Montreal Cognitive Assessment (MoCA). A MoCA score of > 22 was desired to assess the eligibility of all the participants (Carson et al., 2018). Further, a general health questionnaire was used, which revealed that all participants were in good physical and mental health. Additional exclusion criteria included a history of neurological disorders, addiction, stroke, smoking or any contraindications for Magnetic Resonance Imaging (MRI). All participants provided their written consent before any data was collected. The protocol was approved by the local ethical committee of KU Leuven/UZ Leuven, Belgium (project number: S66650). Because the current task was part of the larger data collection to investigate effects of aging on (cerebellar) motor function, this task was performed during a 2.5-hour test session in combination with three other cerebellar motor task, a cognitive task, a fine motor task, and a physical fitness task. The pure-elbow motion task was performed at the beginning of this session and lasted approximately 30 minutes. Only the data of this task was included here.

### 2.2. Experimental Setup

#### Robotic device

We used the KINARM exoskeleton (Kinarm, BKIN Technologies Ltd., Kingston, ON, Canada) to assess kinematics of the upper limbs. The participants were seated in the exoskeleton with their upper limbs supported through three arm rests (upper arm, lower arm and hand). These supports were individually adjusted to enable smooth shoulder and elbow rotations in the horizontal plane, with the shoulders placed in 85° abduction thanks to the height-adjustable chair. A black screen was used to block the view of participants’ hands and arms. In an augmented reality system, a white dot represented the position of the hand.

#### Muscle activity

During the experiment, we collected muscle activity from the right arm using wireless surface electromyography (sEMG) electrodes (Delsys, Trigno Avanti Sensors, Boston, MA, USA). Before placing the EMG electrodes on the skin surface overlying the belly of the muscle, the skin was cleaned with rubbing alcohol, and the contacts were coated with conductive gel. If needed, the skin was also shaved before placing the electrodes. Rectangular electrodes 37 x 27 x 13 mm were attached parallel to the orientation of the muscle fibers. We recorded two muscles that move the shoulder joint (m. pectoralis major clavicular head, shoulder flexor; m. posterior deltoid, shoulder extensor) and two muscles that move the elbow joint (m. biceps long head, elbow flexor; m. triceps lateral head, elbow extensor). EMG data, recorded with the Delsys software, were synchronized with the KINARM embedded software (Dexterit-E, version 3.9), and saved in separate files for each trial of the task.

### 2.3. Pure-elbow motion task

Participants performed 30° pure-elbow movements, starting in three possible joint orientations (**Figure 1**). The start position of the shoulder joint was always at 60° (external angle). Pure-elbow extension movements were performed either from 105° (position A) to 75° (position B) or from 75° to 45° (position C), and pure-elbow flexion movements were performed from either 45° to 75° or from 75° to 105° (elbow joint external angles). The goal target was positioned in such a way that no movement in the shoulder joint was needed to reach from the start position to the goal target. The start position of the next trial was always the same as the position of the goal target of the previous trial (except for the first trial). Participants were instructed to perform a pure rotation of the forearm around the elbow and to keep their shoulder joint as stable as possible while reaching to the goal target. Furthermore, participants were instructed to reach through the goal target and to stop their movement of the forearm as close as possible to this target. When the hand cursor remained for 1– 1.2s at the start position, a goal target appeared. While reaching to this goal target, the hand feedback cursor disappeared until the start position for the next trial appeared (see Maeda et al., 2017). Visual feedback of the hand position during the reaching movement was eliminated to isolate feedforward control mechanisms underlying coordination of the pure-elbow movement. Feedback about movement duration (i.e., the time from leaving the start position to reaching the target) was provided after each trial using a color code. If the actual movement duration fell within the desired movement duration interval (see below), the goal target turned green. The goal target turned blue or orange if the actual movement duration was too long or too short, respectively.

**Figure 1:**
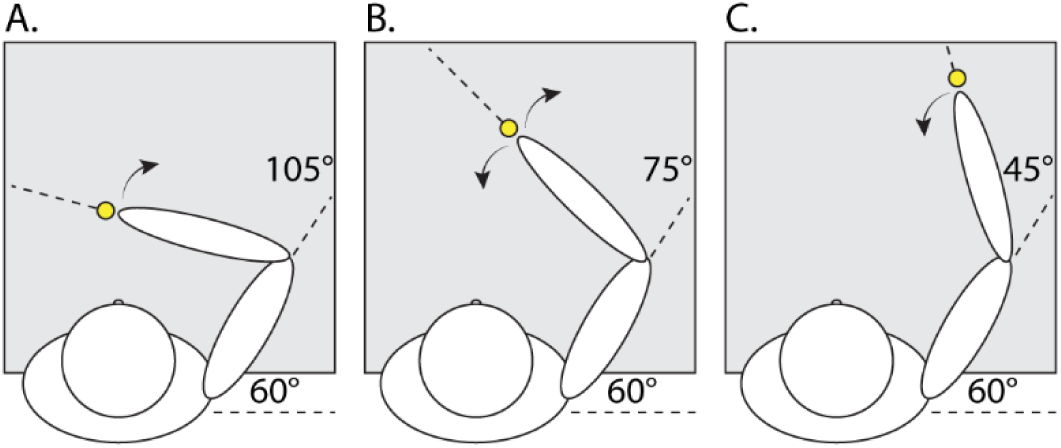
The three possible elbow-shoulder joint configurations at the start of the pure-elbow movement. The shoulder is always positioned at 60° (external angle). Elbow extension movements were made starting from position A or B. Elbow flexion movements were made starting from position B or C.

Participants performed the task under two different speed conditions: a slow and fast condition. Requirements for movement duration were between 250 – 300ms and between 150 – 200ms for the slow and fast condition, respectively. The order of the slow and fast movement speed condition was randomized across participants. Before starting the experimental trials, familiarization trials were provided to inform the participants about the task and the desired movement speed. Participants received 24 and 12 practice trials for the first and second movement speed condition, respectively. Only if the participant could not move the forearm with the desired movement speed for any trial, the familiarization trials were repeated. Participants performed a total of 120 trials in each speed condition (i.e., 30 trials per movement direction and starting position).

Before the task began, normalization trials were performed (see Pruszynski et al., 2008). In these trials, participants were instructed to move the hand cursor to a yellow circle (2 cm diameter), with the shoulder and elbow positioned at 60° and 75°, respectively. During the normalization trials, the exoskeleton gradually applied a torque to either the elbow or shoulder joint, which plateaued at a constant torque of 2 N·m for 4 seconds. These torques included flexion and extension of each joint. Participants were instructed to counter these torques and to maintain the hand cursor positioned in the circle as much as possible. Each participant completed three normalization trials in each condition in a random order. Except for 25 older and 36 young participants, who only completed one normalization trial in each condition. All the experimental conditions described were programmed in MATLAB-Simulink (MathWorks, Natick, MA).

### 2.4. Data analysis

***Joint kinematics* -** Joint kinematics (i.e., hand position, elbow and shoulder angles, angular velocity and angular acceleration) were sampled at 1,000 Hz. Movement onset and end were defined when 5% of peak angular velocity of the elbow joint was reached. All trials were included in the analysis, only trials where participants started and/or completed the movement in the wrong direction were excluded (we used 97.82% of all trials in subsequent analyses). To evaluate shoulder movement during the pure-elbow motion task, we calculated three measures of shoulder displacement based on the angle deviation of the shoulder relative to its start position. These measures were assessed from movement onset until the end of the movement. For each trial, we calculated 1) the total absolute area under the curve, 2) the shoulder-angle deviation at +100ms relative to movement onset, and 3) the deviation angle deviation at the end of the movement. These measures of shoulder stability evaluate (1) the total movement of the shoulder during the elbow movement, (2) the anticipatory shoulder movements, and (3) the accuracy of the shoulder position while reaching the goal target. To assess the accuracy of the performance of the pure-elbow motion task, we additionally investigated the reaching error at the end of the movement. For each trial, we calculated the Euclidean distance in 2D space between the location of the goal target and the position of the hand when the elbow moved 30 degrees toward the target. For all measures, the mean of all trials going in the flexion or extension direction was computed for each participant.

The elbow’s initial configuration and the speed of rotation during the elbow movement will determine the amplitude of the interaction torque that arises at the shoulder. Therefore, we calculated the net interaction torque (𝑇_𝑠_) in the shoulder that depends on the motion of the lower arm for each good trial (see Gribble and Ostry 1999):

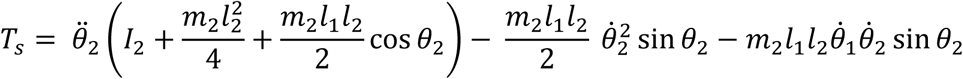

𝜃_1_ and 𝜃_2_ are shoulder and elbow joint angles, 𝜃̇_2_ and 𝜃̈_2_ are elbow velocity and elbow acceleration. 𝐼_2_is the moment of inertia of the lower arm about its center of mass, 𝑚_2_ is the mass of the lower arm, and 𝑙_1_ and 𝑙_2_ are the lengths of the upper and lower arm.

***Muscle activity*** - EMG signals were sampled at 1,000 Hz, bandpass filtered (20-100 Hz, 2-pass, 2^nd^-order Butterworth), full-wave rectified and normalized such that a value of 1 represents a given muscle sample’s mean activity during movements to counter a 2 N·m torque (see Pruszynski et al., 2008). First, we calculated separately for each speed condition the mean muscle activity for all good trials either performed in the flexion or extension direction. Then, based on the mean muscle activity of all these trials in either the flexion or extension direction, we calculated the EMG outcome measures.

We assessed (1) the onset of muscle activity based on the first phasic EMG burst (i.e., EMG onset). To determine the timing of this burst, we calculated baseline EMG activity over a fixed time window from −400ms to −300ms relative to movement onset. Then, we scored the first phasic EMG burst based on the time at which the EMG signal rose 3 SD above the mean baseline EMG activity and remained above that level for at least 50ms. Additionally, we verified that this method to determine the first EMG burst, based on the mean EMG activity of all trials going in the same direction, was correlated with the method where the first EMG burst is calculated from individual trials (p<0.001; r=0.23), with slightly earlier EMG onset detected using the mean-based method. We therefore based subsequent analyses on the mean of all trials in the same movement direction, as this reduces the impact of individual trials that fail to meet the criteria to detect the first EMG burst, providing a more robust estimation of EMG onset per participant.

Further, we assessed (2) the amplitude of muscle activity. First, we subtracted the mean baseline EMG activity from the EMG trace. Then, we calculated the area under the curve with the *cumtrapz* function in MATLAB (Mathworks, Natick, MA, version 2024b) over a fixed time window of −100ms to +100ms relative to movement onset. This time window was chosen to capture the burst of EMG activity for the agonist muscle at the shoulder and elbow.

***Data/Participant exclusion*** - Before starting data analysis of the EMG data, several participants from each age group were excluded because the data were not saved properly. Specifically, the data for the m. deltoid was not saved properly for 6 young adults and for 14 older adults, resulting in exclusion of these participants from all trials with extension movements. Similarly, 2 older-old adults were excluded from all trials with extension movements due to missing data for the m. triceps. Additionally, 1 young adult and 1 older adult were excluded for all trials with flexion movements due to missing data of the m. pectoralis and m. biceps, respectively. After identifying the first phasic EMG burst, additional participants were excluded because, in some cases, the first EMG burst could not be detected for both agonist muscles of the shoulder and elbow within the −300ms to +100ms time window relative to movement onset. See **Table 1** for an overview of the total number of participants with valid EMG data in each age group, movement direction, and speed condition.

**Table 1:** n = the number of participants that have valid EMG data for all the EMG outcome measures in each age group, movement direction, and speed condition. For all the included participants we were able to determine the first phasic EMG burst (i.e., EMG onset) for both the m. biceps and m. pectoralis for trials with flexion movements, or for both the m. triceps and m. deltoid for trials with extension movements (i.e., agonist muscles for each movement direction).

| Age group | Total (n) | Properly saved EMG data (n) | Usable data for all EMG outcome measures (n) |  | % of properly saved data |
| --- | --- | --- | --- | --- | --- |
| 20 - 35<br>Young | 50 | Flexion 49 | Slow | 29 | 59,2% |
|  |  |  | Fast | 40 | 81,6% |
|  |  | Extension 44 | Slow | 32 | 72,2% |
|  |  |  | Fast | 34 | 77,3% |
| 55 - 70<br>Older | 80 | Flexion 78 | Slow | 42 | 53,8% |
|  |  |  | Fast | 63 | 80,8% |
|  |  | Extension 65 | Slow | 51 | 78,5% |
|  |  |  | Fast | 52 | 80,0% |
| 80+<br>Older Old | 30 | Flexion 30 | Slow | 15 | 50,0% |
|  |  |  | Fast | 20 | 66,7% |
|  |  | Extension 28 | Slow | 24 | 85,7% |
|  |  |  | Fast | 26 | 92,9% |

### 2.5. Statistical analysis

Data processing and statistical analyses were performed using MATLAB (Mathworks, Natick, MA, version 2024b). We performed a combination of statistical tests, including repeated measures ANOVA, mixed-design ANOVA, and one-way ANOVA analyses, using the MATLAB functions *fitrm*, *ranova*, and *anova1*, respectively. For the statistical analysis of the timing of the first EMG burst we started with a repeated measures ANOVA with muscle group as within within-factor to check whether the shoulder muscle was activated before the elbow muscle. A separate analysis was conducted for each movement direction and movement speed (flexion-slow: n=86; flexion-fast: n=123; extension-slow: n=106; extension-fast: n=112). Secondly, we performed a mixed-design ANOVA analysis with speed as a within-subject factor and age group as a between-subject factor. In this analysis, the dependent variable was the difference between the timing of the first EMG burst of the shoulder muscle and the timing of the first EMG burst of the elbow muscle. This analysis was performed separately for flexion (n=86) and extension (n=106) movements. Finally, we verified the effect of age group by conducting a one-way ANOVA where only data of the fast speed condition was included, increasing our sample size and statistical power (flexion: n=123; extension: n=112).

For the analysis of the magnitude in shoulder muscle activation, we started with an ANCOVA analysis with age group as a between-subject factor and the maximal interaction torque in the expected direction (𝑇_𝑠_), as a covariate. The effect of age was checked separately for each movement direction and speed condition (flexion-slow: n=86; flexion-fast: n=123; extension-slow: n=106; extension-fast: n=112). Further, a mixed-design ANOVA analysis was performed with speed as a within-subject factor and age group as a between-subject factor in both movement directions (flexion: n=86; extension: n=106). Additionally, we performed and ANCOVA analysis with age group as a between-subject factor and the maximal interaction torque (𝑇_𝑠_), as a covariate. In this analysis, the dependent variable was the difference in the magnitude of shoulder muscle activation between the fast and slow speed condition.

For all the kinematic outcome measures we conducted a mixed-design ANOVA with speed and movement direction as a within-subject factors, and age group as a between-subject factor (n=160). Post-hoc pairwise comparisons between factor levels were conducted with adjustment for multiple comparisons using the multcompare function in MATLAB (Tukey-Kramer method). All results were considered statistically significant if the corrected P value was <0.05.

## Results

We investigated a multi-joint coordination task in which participants reached with their dominant arm to targets arranged in such a way that it required pure forearm rotation around the elbow. Participants were instructed to stabilize their shoulder joint while only moving their forearm. Successful performance of the task required appropriate anticipatory activation of the shoulder agonist muscle to counteract for the interaction torque at the shoulder resulting from forearm motion.

### Feedforward anticipatory mechanism activates the shoulder muscle before the elbow muscle

We tested whether shoulder agonist muscle activation (i.e., m. pectoralis in elbow flexion movements; m. deltoid in elbow extension movements) anticipated the activation of the elbow agonist muscle (i.e., m. biceps in elbow flexion movement; m. triceps in elbow extension movements). **Figure 2A** illustrates the onset timing of the shoulder relative to the elbow muscle, presented separately for each age group. Considering the slow speed condition, we found that the onset of the first EMG burst in the shoulder muscle consistently preceded that of the elbow muscle. This result held for both elbow extension (**Figure 2B**; main effect of muscle: F(1,105)=33.1, p<0.001) and elbow flexion movements (**Figure 2C**; main effect of muscle: F(1,85)=25.6, p<0.001). During elbow extension movements, the shoulder muscle was activated 57±17ms before movement onset and the elbow muscle was activated 46±17ms before movement onset. For elbow flexion movements, the shoulder and elbow muscles were activated 56±23ms and 42±17ms before movement onset, respectively. To verify these results, we repeated the analysis using all participants for whom valid EMG data were available in the fast speed condition, increasing the sample size and statistical power (see **Table 1**). We obtained very similar results to those observed in the slow speed condition for both elbow extension and flexion movements (main effect of muscle: extension: F(1,111)=70.4, p<0.001, shoulder=61±18ms, elbow=47±17ms; flexion: F(1,122)=11.4, p<0.001, shoulder=49±21ms, elbow=42±18ms), supporting our previous findings.

**Figure 2:**
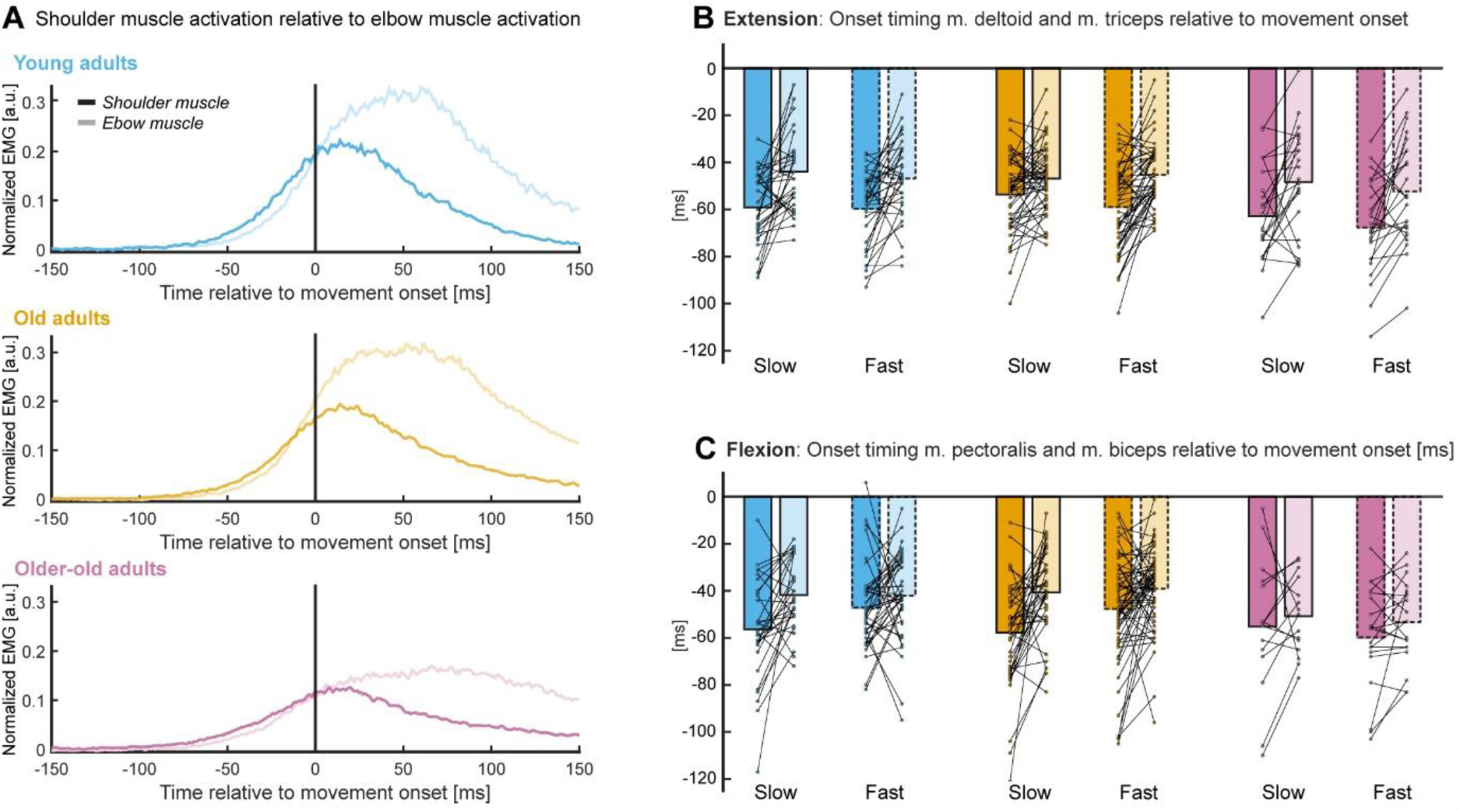
**A**) Illustration of the EMG trace of the shoulder muscle (base color) relative to the elbow muscle (lighter shade of the same base color). The data of the young adults is shown in blue, of the older adults in orange, and of the older-old in purple. The shown data represents the mean EMG trace of all participants in each age group for which we were able to identify the first EMG burst for both the shoulder (m. deltoid) and elbow muscle (m. triceps), in the fast speed condition for elbow extension movements. **B**) Shows the timing of the first EMG burst for the m. pectoralis (base color) and the m. biceps (lighter shade of the same base color) relative to movement onset in both the slow (solid line around bars) and fast (dashed line around bars) movement speed condition. The data is based on all good trials going in the flexion direction. **C**) Shows the timing of the first EMG burst of the m. deltoid (base color) and the m. triceps (lighter shade of the same base color) relative to movement onset in both the slow (solid line around bars) and fast (dashed line around bars) movement speed condition. Data is based on all good trials going in the extension direction. **Panel B and C**) Each dot is representing the timing of the first EMG burst of an individual participant and the height of the bars is representing the mean of all participants in a specific condition. The dots representing the timing of the first EMG burst of the shoulder and elbow muscle of the same participants are connected.

### Ageing does not affect the timing of anticipatory shoulder muscle activation

The results of the previous analysis made clear that we could detect the anticipatory mechanism responsible for the smooth coordination of the shoulder and elbow joints during pure-elbow movements. As illustrated in **Figure 2**, this anticipatory mechanism could be observed for all age groups. Indeed, considering participants with valid data in both speed conditions, we did not find evidence that the relative timing of the shoulder muscle with respect to the elbow muscle changed with age (main effect of age group: extension: F(2,103)=0.76, p=0.47 flexion: F(2,83)=0.75, p=0.47). For elbow extension movements, the shoulder muscle was activated 13±19ms, 10±16ms, and 14±20ms before the activation of the elbow muscle for the young, older and older-old adults, respectively. For elbow flexion, the onset of the anticipatory activation of the shoulder relative to the elbow muscle was 11±21ms, 13±25ms, and 5±21ms for the young, older, and older-old adults, respectively. This absence of significance could be due to the absence of an effect or to limited power. Therefore, the analysis was repeated with all participants with reliable data in the fast speed condition, increasing the sample size to maximize the statistical power and the ability to detect age-related differences (see **Table 1**). These results confirmed the absence of significant differences in the onset timing of shoulder muscle activity relative to onset timing of the elbow muscle activity between all three age groups (main effect of age group: extension: F(2,109)=0.16, p=0.85, YA=13±18ms, OA=14±17ms, OOA=15±19ms; flexion: F(2,120)=0.26, p=0.77, YA=5±25ms, OA=9±24ms, OOA=7±18ms). This means that participants from all age groups exhibited similar anticipatory strategies to compensate for the interaction torque around the shoulder resulting from pure-elbow movement.

### Aging does not affect anticipatory timing under increased motor demands

While age did not affect the relative shoulder-elbow muscle onset timing, we further investigated whether movement speed could challenge this feedforward anticipatory mechanism that drives anticipatory shoulder activation preceding elbow activation. We found that the relative timing between the onset of shoulder muscle activity and the onset of elbow muscle activity was similar for both the slow and fast speed condition (main effect of instructed speed: **Figure 2B**; extension: F(1,103)=0.035, p=0.85; **Figure 2C**; flexion: F(1,83)=3.6, p=0.06). For elbow extension movements, the onset of shoulder muscle activity preceded that of the elbow muscle activity with 12±19ms and 12±17ms in the slow and fast speed condition, respectively. During elbow flexion movements, the onset of shoulder muscle activity preceded that of the elbow muscle activity with 12±25ms and 7±20ms in the slow and fast speed condition, respectively.

Interestingly, across all age groups, no differences between the slow and fast speed condition were observed in the modulation of the shoulder-elbow muscle onset activation timing for elbow flexion movements (interaction between instructed speed and age group: F(2,83)=1.41, p=0.25). However, for elbow extension movements this modulation of shoulder-elbow muscle onset activation timing across speed conditions was different between age groups (interaction between instructed speed and age group: F(2,103)=4.11, p=0.02). Remarkably, older adults appeared to cope with the increased demands on the motor system in the fast speed condition more effectively than younger adults (YA vs. OA: p=0.02). Specifically, older adults initiated the anticipatory shoulder muscle activation, relative to the elbow muscle activation, earlier in the fast speed condition compared to the slow speed condition. In contrast, both young and older-old adults showed a slight tendency to delay the timing of shoulder muscle activation relative to elbow muscle activation when movement speed increased. However, no additional differences between age groups were observed in the modulation of shoulder-elbow muscle activation timing (YA vs. OOA: p=0.77, OA vs. OOA: p=0.19). These results suggest an absence of systematic age-related impairments in the timing of anticipatory shoulder activation, even under more challenging motor demands.

### Ageing does not affect the magnitude of anticipatory shoulder muscle activation

In addition to preserved anticipatory timing, effective feedforward control also requires that the magnitude of shoulder muscle activation is properly scaled with the interaction torque to ensure shoulder stabilization during elbow motion. Therefore, we first examined whether the angular velocity of the elbow and the resulting interaction torque differed between age groups, before comparing shoulder activation magnitude. Differences between age groups in elbow angular velocity (**Figure 3C** & **Figure 3G**; main effect of age group: F(2,156)=3.42, p=0.035) and interaction torque (**Figure 3D** & **Figure 3H**; main effect of age group: F(2,156)=4.83, p=0.009) were quite clear in our sample. Across both speed conditions, the older-old adults performed the task with the lowest angular velocity of the elbow (157±38deg/s) coupled with the lowest interaction torque at the shoulder (0.59±0.41N·m). Older adults performed the task in general slightly faster (164±38deg/s) than the young adults (162±34deg/s), which resulted in the highest interaction torque at the shoulder for the older adults (OA=0.78±0.53N·m, YA=0.69±0.45N·m). Post-hoc comparisons indicated that elbow angular velocity and the interaction torque were significantly higher for the older adults compared to the older-old adults (elbow angular velocity: OOA vs. OA: p=0.024; interaction torque shoulder: OOA vs. OA: p=0.007). However, we did not find differences between the young adults and both older age groups (elbow angular velocity: YA vs. OA: p=0.756, YA vs. OOA: p=0.148; interaction torque shoulder: YA vs. OA: p=0.224, YA vs. OOA: p=0.281).

**Figure 3:**
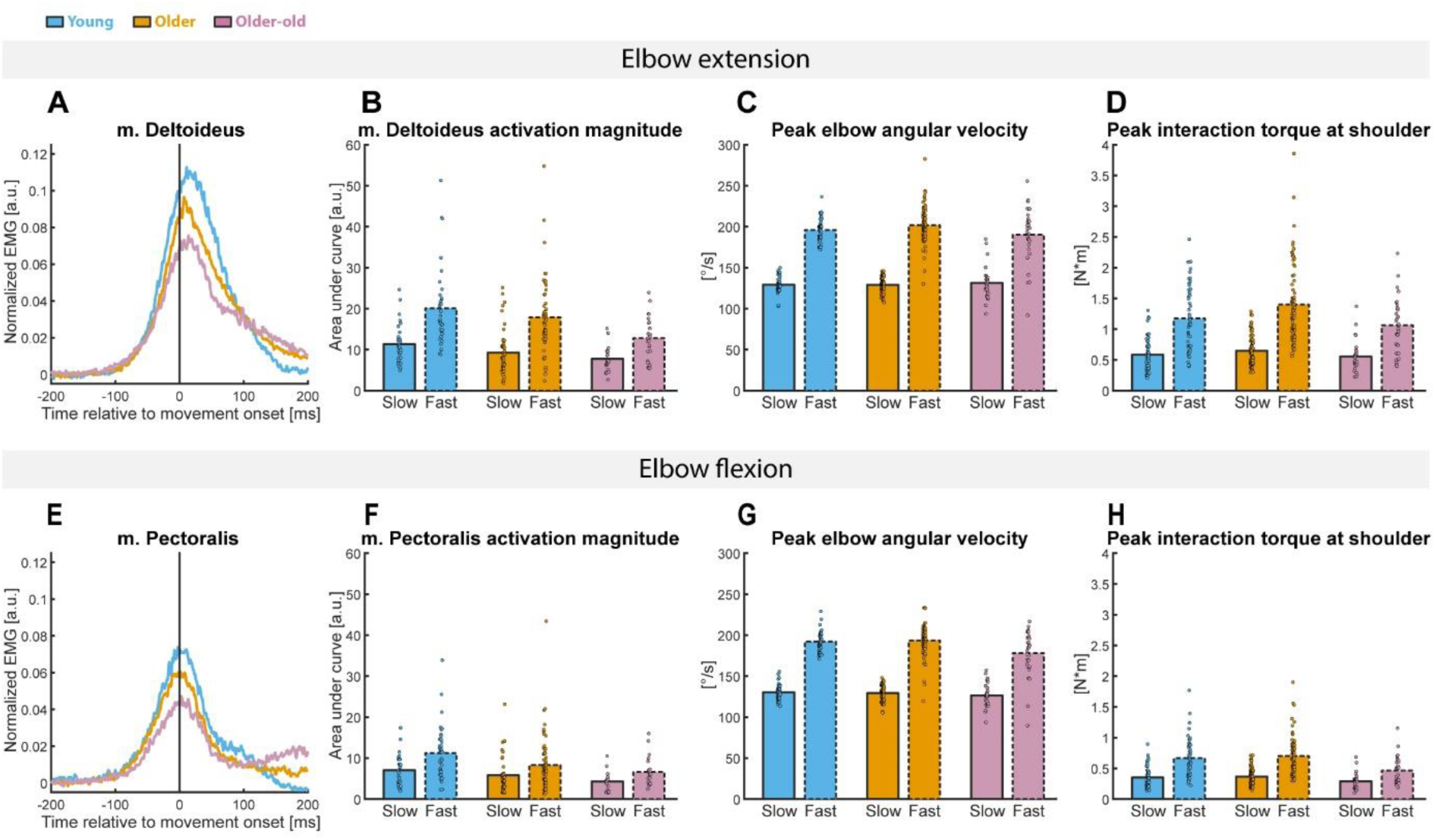
**A**) Mean EMG trace of the m. deltoid for each age group during elbow extension movements (slow condition only). **E**) Mean EMG trace of the m. pectoralis for each age group during elbow flexion movements (slow condition only). **B**) Activation magnitude of m. deltoid, based on the area under the curve of the EMG trace in a time window of −100 to +100ms relative to movement onset. **F**) Activation magnitude of m. pectoralis, based on the area under the curve of the EMG trace in a time window of −100 to +100ms relative to movement onset.. **C**) Peak angular velocity of the elbow joint during elbow extension movements. **G**) Peak angular velocity of the elbow joint during elbow flexion movements. **D**) Peak interaction torque at the shoulder during elbow extension movements. This variable is used as a covariate in the statistical analysis for age related effects on the activation magnitude of the m. deltoid. **H**) Peak interaction torque at the shoulder during elbow flexion movements. This variable is used as a covariate in the statistical analysis for age related effects on the activation magnitude of the m. pectoralis. **Panel A, B, E & F**) These values are not corrected for the age-related differences in magnitude of the interaction torque (i.e. covariate in statistical analysis). Therefore, visual representation of age-related differences will be mainly caused by age-related differences in the magnitude of the interaction torque. **Panel B, C, D, F, G & H**) Each dot is representing the mean value of all trials going in the same movement direction of an individual participant. The height of the bars represents the mean muscle activity of all participants of the corresponding age group. **All Panels**: The blue color represents the data of the young adults, the orange color represents the data of the older adults and the purple color represents the data of the older-old (80+) adults.

To account for these differences in interaction torque across age groups, the interaction torque was included as a covariate in the statistical analyses assessing the effects of aging on shoulder muscle activation magnitude. Considering the slow speed condition, we found that the modulation of the magnitude of shoulder activity was very similar across all age groups (**Figure 3A** & **Figure 3E**; main effect of age group: extension: F(2,100)=0.054, p=0.95, YA=11.3±4.6a.u., OA=11±5.5a.u., OOA=11.4±4.4a.u.; flexion: F(2,81)=0.18, p=0.83, YA=7.1±3.7a.u., OA=6.9±3.7a.u., OOA=6.4±3a.u.). The absence of age-related differences in the magnitude of shoulder muscle activity during the task was confirmed with the data from the fast speed condition, for which power is maximized. In the fast condition, participants from all age groups exhibited very similar magnitudes of shoulder muscle activation when accounting for differences in interaction torque (main effect of age group: extension: F(2,106)=0.059, p=0.94; YA=20.1±9.8a.u., OA=20.6±10.5a.u., OOA=19.1±8.9a.u.; flexion: F(2,118)=0.69, p=0.51; YA=11.3±6.3a.u., OA=10.5±5.9a.u., OOA=9.3±3.9a.u.). This suggests that both the older and older-old adults were able to activate their agonist shoulder muscle to a similar extent as the young adults to effectively compensate for the subsequent interaction torque at the shoulder.

### Aging does not affect anticipatory activation magnitude under increased motor demands

Increasing movement speed increases the challenge for the motor system to appropriate scale shoulder muscle activation to the interaction torque. We found that the peak angular velocity of the elbow significantly increased with the instructed speed (**Figure 3C** & **Figure 3G**; main effect of instructed speed: F(1,156)=1564, p<0.001, slow=129±11deg/s, fast=194±21deg/s), which in turn led to a corresponding increase in the interaction torque at the shoulder (**Figure 3D** & **Figure 3H**; main effect of instructed speed: F(1,156)=408, p<0.001, slow=0.48±0.25N·m, fast=0.96±0.55N·m). Importantly, across age groups, this increase in interaction torque was matched by a proportional increase in the magnitude of anticipatory shoulder agonist activation, demonstrating that the motor system effectively compensated for the higher mechanical demands when elbow velocity was increased (main effect of instructed speed: **Figure 3B**; extension: F(1,101)=373, p<0.001, slow=11.2±5.0a.u., fast=20.0±10.0a.u.; **Figure 3F**; flexion: F(1,82)=196, p<0.001, slow=6.9±3.6a.u., fast=10.8±5.9a.u.).

Notably, this increase in shoulder muscle activation, when elbow velocity was increased, differed across age groups (**Figure 3B** & **Figure 3F**; interaction between instructed speed and age group: extension: F(2,101)=6.25, p=0.003; flexion: F(2,82)=3.75, p=0.027). For both extension and flexion movements, the older-old adults exhibited a smaller modulation of the magnitude of the shoulder muscle activity with instructed speed, compared to younger and older adults (changes in shoulder muscle activity across speed conditions: extension: YA=9.27±5.87a.u., OA=9.53±11a.u., OOA=6.81±10.5a.u.; flexion: YA=4.76±3.55a.u., OA=4.02±7.28a.u., OOA=2.28±3.6a.u.). However, none of the between-group comparisons were statistically significant (extension: YA vs. OA: p=0.99, YA vs. OOA: p=0.61, OA vs. OOA: p=0.49; flexion: YA vs. OA: p=0.85, YA vs. OOA: p=0.36, OA vs. OOA: p=0.57).

However, these results are inconclusive as these age-related differences in the modulation of the shoulder EMG activity with speed could be due to an age effect or to an age-related difference in the changes in interaction torque across speed conditions.

Indeed, participants from different age groups modulated the elbow angular velocity and the corresponding interaction torque differently with the instructed speed (interaction between age group and instructed speed: **Figure 3C** & **Figure 3G**; elbow angular velocity: F(2,156)=5.70, p=0.004; **Figure 3D** & **Figure 3H**; interaction torque (F(2,156) = 6.96, p=0.001). While the

maximum angular velocity of the elbow was very similar across all age groups in the slow speed condition (YA=129.9deg/s, OA=129.3deg/s, OOA=129.1deg/s), older-old adults increased their elbow angular velocity less with instructed speed than young adults (OOA vs. YA: p=0.014) and older adults (OOA vs. OA: p<0.001) (change in elbow angular velocity across speed conditions: YA=64.3±10.4deg/s, OA=68.7±19.6deg/s, OOA=55.3±29.7deg/s). This smaller increase in elbow angular velocity resulted in smaller increase in interaction torque across speed conditions for the older-old adults, compared to the young (OOA vs. YA: p=0.13) and older adults (OOA vs. OA: p=0.004) (changes in interaction torque across speed conditions: YA=0.45±0.28N·m, OA=0.55±0.4N·m, OOA=0.34±0.32N·m). The latter result explains why older-old adults exhibit a smaller increase in shoulder EMG activity across instructed speed conditions compared to younger and older ones: a smaller change in interaction torque requires a smaller change in shoulder muscle activity. This interpretation is confirmed by the absence of interaction effect when the interaction torque is added to the statistical model as a covariate (interaction between age group and instructed speed: extension: F(2,100)=0.12, p=0.887; flexion: (F(2,81)=0.92, p=0.403). Together, these results suggest that all participants, independently of their age group, exhibited similar activation magnitudes of the shoulder muscle during the task, and modulated it appropriately under increased motor demands when elbow velocity increased.

### Preserved anticipatory mechanisms are not sufficient to prevent age-related differences in accuracy by the end of the movement

On the muscle level we could not identify any age-related differences, indicating that the anticipatory mechanisms remain intact during healthy aging. However, such intact anticipatory behavior is not enough to guarantee perfect shoulder stability during the whole elbow movement. Indeed, the total deviation angle, measured from movement onset to movement offset, increased with increasing age (**Figure 4B** & **Figure 4F**; main effect of age group: F(2,157)=8.5, p<0.001, YA=1.52±0.84deg/s, OA=1.89±0.91deg/s, OOA=2.22±1.55deg/s). The older and the older-old adults exhibit larger upper arm motion compared to the young adults (YA vs. OA: p=0.018; YA vs. OOA: p<0.001). This was also larger in older-old adults than older adults, but this difference failed to reach significance (p=0.098). Increasing movement speed increased the magnitude of the shoulder movement (slow=1.74±1.07deg/s, fast=1.93±1.06deg/s, F(1,157)=15.79, p<0.001) and this effect was consistent across all age groups (interaction between age group and instructed speed: F(2,157)=0.83, p=0.44).

**Figure 4:**
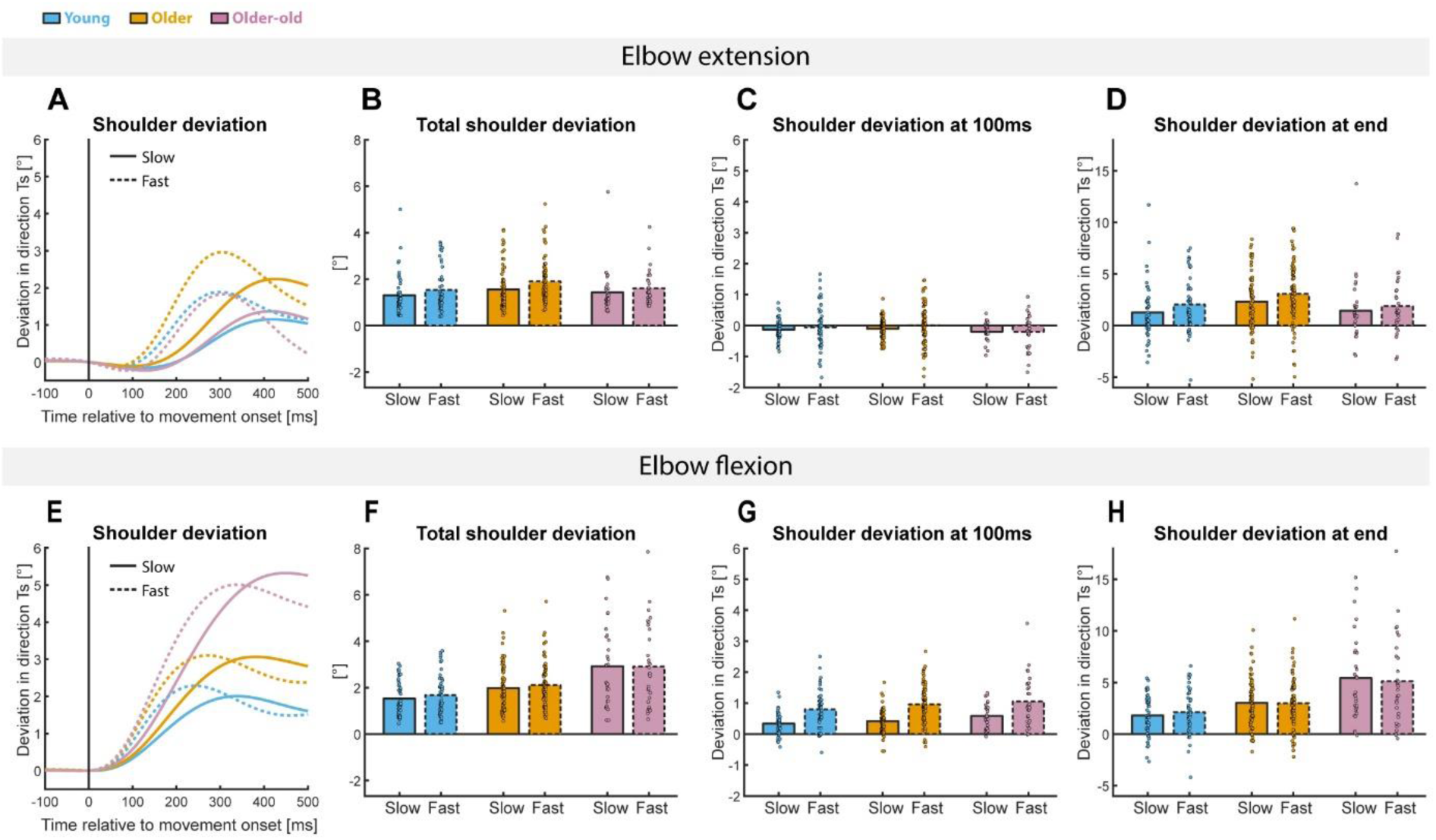
Panel **A & E**) Deviation angle, in the expected direction of the interaction torque, relative to its start position (i.e. position at the time of movement onset). Panel **B & F**) Total deviation angle from movement onset until movement end (based on elbow velocity). **Panel C & D**) Deviation of the shoulder, in the expected direction of the interaction torque, relative to its start position at 100ms after movement onset. Negative values indicate deviation of the shoulder in the opposite direction of the interaction torque at the shoulder. **Panel D & H**) Deviation of the shoulder, in the expected direction of the interaction torque, relative to its start position at the end of the movement (based on elbow velocity). **All panels**) The slow and fast speed condition are indicated with solid and dashed lines, respectively. The blue color represents the young adults, the orange color the old adults, and the purple color the older-old adults.

Given the observed decline in shoulder stability with older age, we further investigated whether these age-related differences were already present during the initial phase of the elbow movement. Across all age groups, deviation angle within the first 100ms of the movement was very minimal, consistently remaining below 0.5° (**Figure 4C** & **Figure 4G**; main effect of age group: F(2,157)=0.96, p=0.39, YA=0.24±0.64°, OA=0.32±0.65°, OOA=0.31±0.76°). This suggests that in the initial phase of the elbow movement, older and older-old adults were still able to compensate for the interaction torque arising at the shoulder to the same extent as the young adults. Increasing movement speed did significantly cause more deviation of the shoulder position in the initial phase (main effect of instructed speed: F(1,157)=96.15, p<0.001, slow=0.15±0.44°, fast=0.44±0.81°). Nevertheless, the effect of increasing movement speed on shoulder stability in the initial phase of the elbow movement was similar across all age groups (interaction between age group and instructed speed: F(2,157)=1.01, p=0.37).

However, the effect of the interaction torque on the shoulder stability accumulated over time such that the amount of deviation angle increased during the movement and this effect was more pronounced in older adults. Indeed, the amount of deviation angle at the end of the elbow movement varied across age groups (**Figure 4D** & **Figure 4H**; F(2,157)=7.17, p=0.001, YA=1.82±2.36°, OA=2.85±2.6°, OOA=3.5±4°). The deviation angle at the end of the movement of the young adults was significantly smaller compared to that of the older adults (p=0.014) and to that of older-old adults (p=0.001). Older-old adults also deviated their shoulder more compared to older adults, but this difference was not significant (p=0.44). Furthermore, participants deviated their shoulder more if movement speed was increased (main effect of instructed speed: F(1,157)=7.56, p=0.007, slow=2.5±2.9°, fast=2.8±2.9°), but the effect of speed was similar across all age groups (interaction between age group and instructed speed: F(2,157)=1.2, p=0.31). These results suggest that the intact anticipatory behavior at the muscle level will mainly support shoulder stability at the initial phase of the elbow movement, but not at the end of the movement.

The lack of full shoulder stabilization in the groups of older participants is reflected in the accuracy of reaching the endpoint. In the absence of any visual feedback of the hand position during the reach, older adults performed the task less accurately than the younger participants and their hand position was further away from the target position (**Figure 5**; main effect of age group: F(1,157)=1078, p<0.001, YA=1.22±0.58cm, OA=1.45±0.68cm, OOA=1.79±1.14cm).

**Figure 5:**
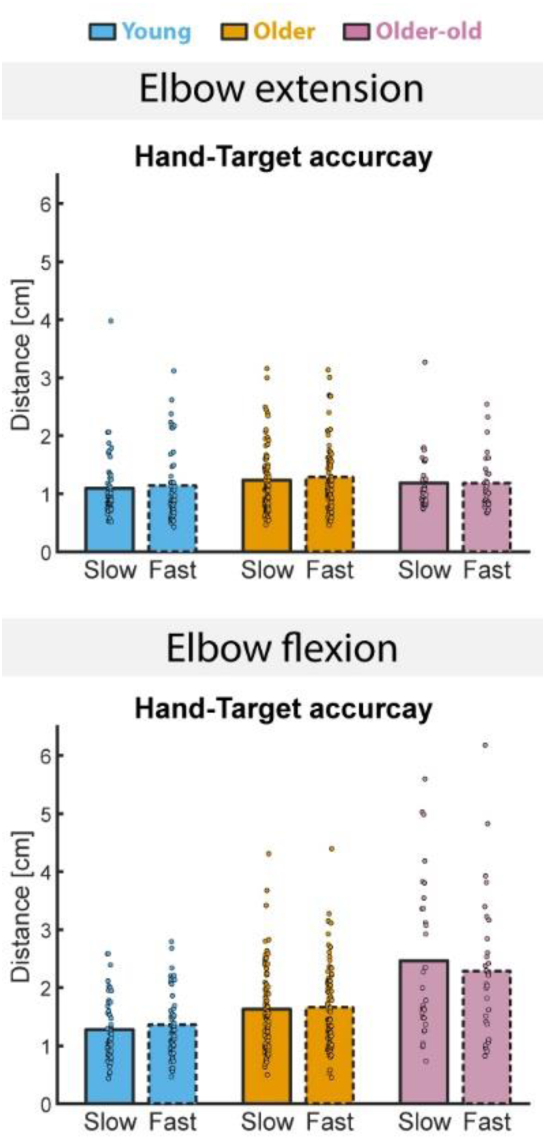
Hand-target accuracy based on the distance between the position of the target and the position of the hand when the elbow moved 30° in the direction of the target. The slow and fast speed condition are indicated with solid and dashed lines, respectively. The blue color represents the young adults, the orange color the old adults, and the purple color the older-old adults.

The endpoint accuracy differed significantly between each age group (YA vs. OA: p=0.038; YA vs. OOA: p<0.001; OA vs. OOA: p=0.009), but there was no evidence that performance was affected by the instructed speed (F(1,157)=0.03, p=0.86, slow=1.43cm, fast=1.46cm). Furthermore, there is no evidence that the reach accuracy is differently affected by instructed speed across age groups (interaction between age group and instructed speed: F(2,157)=1.89, p=0.15). These results suggest that a lack of shoulder stability throughout the elbow movement could affect the accuracy of the general reach performance.

## Discussion

### Summary

This study investigated the effects of healthy aging on the anticipatory control of intersegmental dynamics in the arm. This was investigated during a pure-elbow motion task, which required active stabilization of the shoulder joint. Our results demonstrate that the anticipatory behavior does not deteriorate during healthy aging. All age groups, including young, older and older-old adults, were able to appropriately time their shoulder muscle activation such that it preceded the activation of the elbow muscle, which is necessary to counteract the interaction torque that will arise at the shoulder because of the elbow movement. Furthermore, older and older-old adults were able to scale their activation of the shoulder agonist muscle to the same extent as young adults to counteract the interaction torque at the shoulder. This appropriate timing and scaling of muscle activation resulted in very minimal deviations of the shoulder position within the first 100ms of the elbow movement, across all age groups. This suggests that participants from all age groups could appropriately plan their movements and take into the complex body dynamics. However, an age-related decrease in shoulder stability became evident toward the end of the elbow movement, resulting in decreased reaching accuracy with older age. Taken together, our results demonstrate that while anticipatory neuromuscular control is preserved with healthy aging, it is not sufficient to maintain shoulder stability beyond the feedforward phase of the movement. These findings may suggest that, in healthy aging, feedback control deteriorates while feedforward control does not.

### Preservation of feedforward control during aging

In our study, we did not observe any age-related declines in the anticipatory control at the muscular level, nor in the kinematic parameters during the initial phase (<100ms) of the reaching movement. Therefore, our findings indicate that feedforward control, which enables the nervous system to anticipate and plan movements, remains intact during healthy aging. One defining feature of feedforward control is anticipatory timing, whereby muscles spanning adjacent joints are activated before joint motion begins (Almeida et al., 1995; Galloway & Koshland, 2002; Gottlieb, 1998; Gottlieb et al., 1989; Gribble & Ostry, 1999; Maeda et al., 2017). Consistent with this principle, both shoulder and elbow muscles in our study were activated prior to movement onset, and this anticipatory activation was preserved across all age groups. Moreover, during the pure-elbow motion task, where the shoulder was free to move, we found that shoulder muscle activation consistently preceded elbow muscle activation by approximately 11-14ms. This anticipatory activation pattern ensured shoulder stabilization during the initial phase of the elbow movement (<100ms) across all age groups, despite the interaction torque rising from movement onset and peaking at 100-150ms in our task. Another hallmark of intact feedforward control observed in our study was that the magnitude of shoulder muscle activity was appropriately scaled to elbow muscle velocity and the resulting interaction torque at the shoulder. This aligns with prior evidence that shoulder agonist activity not only precedes elbow activation but also takes intersegmental dynamics into account during self-initiated reaching, (Gribble & Ostry, 1999; Maeda et al., 2017). Crucially, we found this scaling preserved across all age groups. Overall, these findings demonstrate no evidence for age-related effects on feedforward control during a multi-joint coordination task of the upper limbs.

These results are in line with previous studies demonstrating that healthy aging does not reduce the quality of feedforward control, and that older adults retain the capacity to adjust feedforward control mechanisms appropriately to maintain accurate movements (Pai et al., 2003; Sager et al., 2024). Accordingly, intact feedforward control with older age has also been observed in the context of vertical arm movements. In such vertical arm movements, participants modulate their motor command to take advantage of the gravity to decelerate upward movements or to accelerate downward movements (Crevecoeur et al., 2009; Gaveau et al., 2016; Gaveau et al., 2011; Gaveau & Papaxanthis, 2011). The ability to plan a movement while taking gravity into account remains unimpaired with age (Mathieu et al., 2024; Poirier et al., 2020). Importantly, in our study we observed that the resilience of anticipatory planning mechanisms was evident not only in older adults aged 55-70, but also in those over 80 years of age. Our results demonstrate that, even in advanced age, the nervous system can accurately predict and compensate for the arising interaction torque at the shoulder. Together with our results, these findings provide compelling evidence that feedforward control mechanisms supporting multi-joint coordination remain preserved across the lifespan. To our knowledge, our study offers the first empirical evidence that anticipatory motor planning remains resilient in adults aged 80 years and older, highlighting the robustness of feedforward control in the aging nervous system.

### Deterioration of feedback control during aging

Although shoulder stability was preserved during the initial phase of the movement, we observed reduced endpoint accuracy with older age. Because correcting the movement trajectory during the movement depends on sensory feedback about the hand position, this decline in endpoint accuracy suggests an age-related deterioration in feedback control. The deterioration of these feedback mechanisms may result either from noisier proprioceptive input or from deficits in the neural processing of this sensory information. Consistent with the first possibility, Konczak et al. (2012) found increased noise in the incoming sensory signals during movements at older age. In accordance to these results, previous research has shown altered subcortical processing linked to poorer proprioceptive abilities in older adults (Goble et al., 2012). These age-related effects on proprioception could make it more challenging for the nervous system to accurately estimate the hand position and the movement of the limb (Ferlinc et al., 2019; Proske & Gandevia, 2012), which is crucial for ensuring endpoint accuracy.

Interestingly, increased sensory noise in the motor system of older adults could reduce their reliance on sensory feedback, shifting movement control toward predictive mechanisms (Chan et al., 2021; Parthasharathy et al., 2022; Wolpe et al., 2016). In the absence of visual feedback, as in our experiment, it is therefore likely that older adults relied more heavily on predictive mechanisms when reaching to the target (Bosco et al., 2012). Whereas our results demonstrate that feedforward control remained well preserved across all age groups, greater reliance on predictive mechanisms might have masked subtle age-related deficits in feedback corrections. For example, we found age-related deficits in shoulder stabilization at the end of the elbow movement, but these were less evident during extension compared to flexion. During extension, we accordingly observed less deviation in the position of the shoulder in the initial phase (<100ms), when feedforward control is most critical. This could suggest that greater reliance on preserved feedforward mechanisms may partly compensate for the weakened feedback control, thereby supporting endpoint accuracy in multi-joint reaching movements in older adults.

Although noisier proprioceptive input could increase reliance on feedforward control mechanisms, evidence on age-related deterioration of proprioceptive function remains mixed. While age-related declines in proprioceptive function have often been emphasized, recent studies with larger samples suggest that sensory noise may be less pronounced than previously thought (Djajadikarta et al., 2020; Parthasharathy et al., 2022; Van De Plas & Orban de Xivry, 2025; Saenen et al., 2023). Moreover, physically active older adults appear to maintain proprioceptive acuity in the upper limbs better than inactive peers (Adamo et al., 2009; Kitchen & Miall, 2019). Given that our participants were recruited near a sports campus, and consistent with these recent findings, it is likely that our sample exhibited relatively low age-related increases in proprioceptive noise.

While proprioceptive function might be preserved in our sample, the observed age-related deterioration in feedback control could also result from deficits in neural processing of the sensory information. For example, in our task, reduced neural sensitivity to subtle deviations in shoulder position could impair corrective responses or limit fine-tuning of the elbow movement in the final phase. Such deficits may be associated with altered feedback gains that have been observed in older adults (Griffioen & van Dieën, 2020). For example, studies on postural control have indicated that feedback mechanisms in older adults are engaged only when fluctuations are large (Chen et al., 2021). This reduced sensitivity limits the responsiveness of older adults to smaller movement deviations, potentially inducing corrective adjustments at the end of the movement too late (Chen et al., 2021). Consistent with this view, older adults are found to exhibit slower and less accurate corrections than younger adults in goal-directed reaching tasks (Kimura et al., 2015; Sarlegna, 2006). Thus, previous research has proposed that age-related feedback deficits may result from either proprioceptive deterioration or altered neural processing. Although our experimental design does not allow us to disentangle these mechanisms, the reduced endpoint accuracy observed in our task is consistent with age-related declines in feedback control.

### The cerebellum as key brain area underlying age-related resilience of feedforward control

To execute a well-coordinated reaching movement, relying on feedforward control, the cerebellum plays a crucial role in ensuring that predictive motor commands are precisely calibrated (Bastian et al., 1996, 2000, 2006; Bruttini et al., 2015; Marchese et al., 2020; Shadmehr & Krakauer, 2014). For example, Oh et al. (2025) demonstrated that intact cerebellar function is essential for anticipatory motor control during reaching tasks involving coordination between the shoulder and elbow joint. Their study showed that cerebellar patients exhibited deficits in the early-phase of motion planning and that they were unable to appropriately compensate for the interaction torque. Furthermore, Cao et al. (2025) identified that cerebellar patients exhibited a prolonged feedforward delay of approximately 70ms relative to healthy controls during a virtual 3D reaching task. This temporal discoordination was corelated with poorer target tracking performance.

These findings underscore the critical role of the cerebellum in feedforward mechanisms and the importance of its integrity for appropriate anticipatory motor planning. This suggests that this particular cerebellar function remains intact in all age groups from our study. Importantly, we attempted to challenge the motor system by increasing the movement speed, thereby elevating the interaction torque and the demands on predictive control. While older adults are generally more susceptible to performance declines under such task stressors (Lavretsky & Irwin, 2007; Whitson et al., 2016), we found no age-related impairments in anticipatory motor planning, even in participants over 80 years old. Shoulder activity remained well-scaled to the observed interaction torque, effectively stabilizing the shoulder during the initial phase (<100ms) of the elbow movement, regardless of age or task difficulty. Knowing the cerebellum’s crucial role in feedforward control, these findings are consistent with a recent study from our lab that demonstrated preserved cerebellar motor function, across a variety of cerebellar tasks in the same populations (De Witte et al., 2025). Taken together, these findings show that cerebellar motor function is resilient during healthy aging, leading to preserved feedforward control mechanisms.

### Limitations

First, the onset time estimates of the first EMG burst are likely biased, leading to an underestimation of the onset timing differences between the muscles of the shoulder and elbow. We used a technique that estimates muscle onset timing as the time when the EMG signal reached some threshold relative to baseline. Results of this method will be biased toward earlier values for muscles showing greater activation levels, which are the muscles at the elbow joint (i.e., the moving joint) in our experiment. Therefore, this can lead to an underestimation of the real timing difference between the shoulder and elbow muscles. Furthermore, we calculated the onset time of the first EMG burst based on the mean signal of all trials going in the flexion or extension direction, instead of calculating the onset time based on individual trials. This method could cause a shift in the timing of the first EMG burst. However, this shift will equally influence the timing of the muscles of the elbow and shoulder, wherefore it will not influence the timing difference between them.

Second, our experimental design did not allow us to directly isolate feedback responses, which limits the extent to which our findings can be attributed specifically to age-related declines in feedback control. In our task, feedback control is interpreted as the estimation of hand position when reaching the target. In contrast, studies that explicitly examine age-related effects of feedback control typically include perturbations, such as unexpected target displacements during the movement (Kimura et al., 2015; Sarlegna, 2006). Because our design did not involve perturbations or visual feedback during the elbow movements, the evaluation of the feedback system was less direct. Nevertheless, since intact feedback mechanisms are essential to maintain endpoint accuracy in our task, the observed age-related differences still provide meaningful insights into feedback control during healthy aging.

Third, a substantial number of participants were recruited in the vicinity of the sports campus or had connections with students from the Faculty of Movement and Rehabilitation Sciences. Therefore, our sample is likely fitter than the general population, which could possibly minimize or occlude the age-related differences in motor control that might be more apparent in the general population. However, while it is likely that our young adult group (i.e., reference group to investigate effects of aging) is also fitter than average and performs above population averages, this would rather increase the likelihood of detecting age-related differences than obscuring them. Therefore, the consistency of the potential bias across all age groups does not pose any concerns for identifying age-related effects. As such, our findings remain valid and generalizable within the broader context of healthy aging research.

Finally, we observed that older and older-old adults had more difficulty precisely terminating their movement on the target while simultaneously executing the elbow movements at the desired speed, compared to young adults. To execute the movement at the desired speed, older or older-old participants often altered their movement strategy, which frequently resulted in an overshoot of the target. This overshoot likely reflects the increased task demands experienced by older adults, due to both the spatial requirement of accurately stopping the hand on the target and the temporal requirement of simultaneously reaching it at the desired movement speed. It is likely that the dual-task demands imposed greater cognitive load on older participants, leading to reduced accuracy in reaching the goal target (Hassan et al., 2022; Kahya et al., 2022). However, while feedforward control remained unaffected across all age groups, even when older and older-old adults possibly experienced increased task demands due to dual-tasking, this confirms that our experimental paradigm was sufficiently challenging to detect subtle age-related deficits. With respect to feedback control, the dual-task demands may have amplified age-related deficits (Hassan et al., 2022; Kahya et al., 2022), that under less demanding task conditions could otherwise be compensated by cognitive resources. Therefore, the increased task difficulty due to dual tasking does not undermine our conclusions. Instead, it supports our findings that feedback control is more vulnerable to aging, whereas feedforward control mechanisms remain resilient throughout healthy aging.

## Conclusion

In conclusion, we found that the anticipatory feedforward mechanisms, which are critical for compensating for the intersegmental dynamics, appear to be preserved during healthy aging. In contrast, we observed a decline in kinematic stability towards the end of the elbow movement, suggesting that the feedback mechanisms based on sensory feedback may be affected by aging. Future research should aim to deepen our understanding of how aging influences motor control by integrating both feedforward and feedback mechanisms with changes in brain function. For example, functional assessments of brain regions such as the cerebellum, premotor cortex, and the sensorimotor cortex, which are key brain areas in generating and adjusting motor commands, may be crucial to further understand the age-related changes in motor control. Moreover, it is important to explore these processes under varied and more challenging task conditions to assess the resilience of the motor control system with age. This will allow researchers to study the resilience of motor control mechanisms in the aging nervous system.

## Acknowledgement

We thank our master students for helping with data collection. This work was supported by the Fonds voor Wetenschappelijk Onderzoek Vlaanderen (FWO G095121N). No conflicts of interest are declared by the authors.

